# Modulating p38MAPK recalibrates sensitivity of drug-resistant pancreatic adenoductal carcinoma cells toward chemotherapy and correlate with improved outcome in PDAC patients: Experimental and metadata evidence

**DOI:** 10.1101/2025.10.04.680430

**Authors:** Vandana Mehra, Khushi Gandhi, Prasenjit Das, Gayatri Sharma, Monika Bhardwaj, Céline Gongora, Hridayesh Prakash

**Affiliations:** Amity Centre for Translational Research, Amity University, Noida, Uttar Pradesh, India; Amity Institute of Biotechnology, Amity University, Noida, Uttar Pradesh, India; Department of Pathology, All India Institute of Medical Sciences, Delhi, India; IRCM, Montpellier, Inserm, ICM, CNRS, Montpellier, France

**Keywords:** PDAC, chemoresistance, p38MAPK, Gemcitabine, Ralimetinib, Apoptosis, Clinical correlation

## Abstract

Pancreatic cancer is one of the deadliest cancers and has very limited therapeutic options and a dismal prognosis. Among various signaling pathways which are activated during tumor development, hyperactivation of Mitogen-Activated Protein Kinase (MAPK) is responsible for high grade angiogenesis, polarization of Tumor Associated Macrophages, unfolded protein responses and exhaustion of T cells, which together contributes towards this therapeutic resistance. We therefore believe that MAPK targeting is expected to enhance sensitivity of highly resistant PDAC cells toward various cancer directed interventions. In this context, we investigated the impact of modulating p38MAPK on the sensitivity of pancreatic cancer cells towards gemcitabine. Supporting our hypothesis, our results convincely, demonstrated that indeed, p38 inhibition sensitizes both KRAS positive Panc-1 and MiaPaCa2 pancreatic carcinoma cells towards gemcitabine induced death. Interestingly p38MAPK targeting significantly reduced the cell viability, clonogenic potential of these cells and enhanced the early apoptosis. Our in-silico studies, supporting our in vitro data, potentially correlated that that high expression of p38α MAPK14 in PDAC patients is associated with poor prognosis and disease free survival. Deep miming of in silico data further demonstrated that MAPK14, in association with, hypoxia inducible factor-1 alpha and vascular endothelial growth factor signaling pathways promote angiogenic programming of PDAC which render these tumors refractory for cancer directed interventions. Based on our preliminary data, we believe that p38 MAPK based approach is potential approach for changing the faith of PDAC patients toward chemo and immunotherapy and believed to improve PDAC burden effectively in the host.

**Highlights:**

- p38 MAPK inhibition enhances the sensitivity of KRAS+ pancreatic cancer cells for Gemcitabine
- P38MAPK knockdown cells are sensitive for Gemcitabine induced death
- MAPK14 (p38α) is associated with angiogenesis and poor prognosis in pancreatic cancer
- MAPK14 Regulates Pro-Tumorigenic Pathways and immune infiltration in Pancreatic Cancer

## Introduction

Pancreatic adenoductal carcinoma is the 6^th^ major cause of cancer related deaths worldwide with less than 11% 5-year survival rate (≤ 11%) in the PDAC patients (1). The conventional and / or modern therapeutic interventions could not prolong the survival of tumor patients beyond few days to few weeks. This is largely due to high-grade angiogenic reprograming of patients which offers high grade resistance to chemo and / or radiotherapy. The limited availability of advanced diagnostic tools and therapeutic options further exacerbates this burden. Although immunotherapy has gained considerable traction globally, its application in pancreatic cancer still remains in its early stages. The immune-suppressive tumor microenvironment and the inherent resistance mechanisms of pancreatic cancer pose inbuilt challenges that need recalibration. have been employed in various human cancers, including pancreatic carcinomas. Therapeutic efficacy of various tumor-directed interventions remains passive against pancreatic cancer which is primarily due to high grade angiogenesis, endothelial anergy, accumulation of M2 TAMs and Tregs and unfolded proteins responses which altogether render PDAC microenvironment cold or refractory. The oncogene KRAS and the tumor suppressors, TP53, CDKN2A, and SMAD4 are found to be mutated in over 50% of patients, often with multiple mutations occurring simultaneously in the same individual. Of the four predominant mutated genes, KRAS mutations are the most prevalent, occurring in over 90% of PDAC cases (2, 3). The KRAS gene encodes a small GTPase enzyme that acts as a molecular switch, linking membrane-bound growth factor receptors to downstream signaling pathways. KRAS has a G domain, where GTP binding and hydrolysis occur, and a membrane domain for anchoring and activity.

Mutations in KRAS lead to a constitutively active form bound to GTP, which interacts with over 80 downstream effectors, activating key pathways like RAF-MEK-ERK, PI3K-AKT-mTOR, and MAPK. Mutant KRAS drives oncogenic growth by activating transcription factors that promote cell growth, differentiation, survival, inflammation, and stimulates inflammatory networks, including NF-κB, JAK/STAT, AP-1, and CREB, which enhance cytokine and chemokine production (e.g., TNFα, IL-6, CXCLs), fostering interactions between tumor cells and the tumor microenvironment (TME) via autocrine and paracrine signaling (3). Out of these pathways, p38MAPK is the integral part of EGFR2 signalling therefore inhibiting P38MAPK will curtail angiogenesis and growth signals to tumours and predispose them sensitive for death. Gemcitabine is the first line chemotherapy which is currently used for the treatment of pancreatic cancer in upfront setting. Due to increasing resistance of PDAC patients toward Gemcitabine, this has been tried in combination of nab-paclitaxel or FOLFIRINOX. However, drug resistance still remained a major hurdle.

In this context, our previous studies (4, 5) have demonstrated that systemic irradiation of RipTag5 mice bearing human insulinomas with low-dose gamma irradiation, effectively reprogrammed the M2 macrophage-enriched tumor microenvironment and normalized tumor vasculature in a M1 macrophage-dependent manner. Interestingly low-dose gamma irradiation reduced the expression of p38 MAPK proteins within highly invasive, angiogenic and impermissive tumours and late stage TAM populations which play critical role in macrophage polarization and the regulation of insulin receptor sensitivity and factors crucial for the development of pancreatic tumors.

P38MAPK is the central proteins network which drives as strong desmoplastic / stromal activation which is associated with refractory phenotype of PDAC. Additional studies have also shown that inhibition of the p38 MAPK pathway reduces IL-1α production, suppresses tumor cell migration, and attenuates the inflammatory cancer-associated fibroblast (CAF) phenotype, thereby enhancing the response to chemotherapy and improving survival in murine PDAC models (6). In the realm of signaling pathways, the p38 MAPK pathway has been implicated in various malignancies. Existing research highlights its role in inflammation and cancer progression; however, a comprehensive assessment of its potential to modulate the immunogenic response in pancreatic cancer remained elusive. Addressing these gaps could open new avenues for therapeutic interventions in this malignancy. The p38 MAPK pathway expressed across various cell types and can regulate different biological processes (7, 8) including tumor cell proliferation, survival, cellular response to stress and inflammation, making it a pivotal player in cancer progression, and chemoresistance. Recent studies revealed that p38 acts as a pro-survival in epithelial ovarian cancer and a key mediator of chemotherapeutic resistance (9). Similarly, downregulation of p38 MAPK is expected to overcomes drug resistance in oxaliplatin-resistant colorectal cancer cells (10). Apart from its role in chemoresistance, p38α has also been shown to be a novel target for inhibiting the aggressive growth of colon cancer cells (11) and colorectal cancer stem cells in tumorspheres and organoids, partly through its association with the β-catenin signaling pathway (12). Based on these presented evidences, we sought to explore the effects of p38 MAPK inhibition on the chemosensitivity of pancreatic cancer cells.

Following this rolling hypothesis, our results convincely, demonstrated that indeed, p38 inhibition sensitizes both KRAS+ Panc-1 and MiaPaCa-2 pancreatic carcinoma cells towards gemcitabine induced death. Interestingly p38MAPK targeting significantly reduced the cell viability, clonogenic potential of these cells and enhanced the early apoptosis. Our in-silico studies, supporting our in vitro data, potentially correlated that that high expression of p38α (MAPK14) in PDAC patients is associated with poor prognosis and disease free survival. Deep miming of in silico data further demonstrated that MAPK14, in association with, hypoxia inducible factor-1 alpha and vascular endothelial growth factor signaling pathways promote angiogenic programming of PDAC which render these tumors refractory for cancer directed interventions. Based on our preliminary data, we believe that p38 MAPK based approach is potential approach for changing the faith of PDAC patients toward chemo / immunotherapy and believed to improve PDAC burden effectively in the host.

## Results

### p38 MAPK inhibition by Ralimetinib (RLI) enhances the sensitivity of Gemcitabine (Gem) against pancreatic cancer cells

To determine the specific role of p38 MAPK in Gem resistance in human pancreatic cancer cells, we assessed the cytotoxic effects of the p38 MAPK pharmacological inhibitor RLI and the chemotherapeutic drug Gem. The half-maximal inhibitory concentration (IC_50_) of each drug in MIA PaCa-2 pancreatic cancer cells were calculated following treatment for 48 and 72 hours. The IC₅₀ values of Gem were 0.6 µM and 0.4 µM at 48 and 72 hours **(Fig. 1A)**, while those of RLI were 18 µM at 48 hours and 13 µM at 72 hours **(Fig 1B)**, respectively. IC₅₀ is a reference point for single-drug potency and it does not account for drug-drug interactions, therefore it is not suitable for combination studies. We selected 13 µM (RLI) and 0.4 µM (Gem) for combination studies in MIA PaCa-2 cells. To determine the role of p38 MAPK in Gem resistance, in pancreatic cancer cells, MIA PaCa-2 cells were pre-treated with RLI for 24 hours, followed by co-treatment with Gem for an additional 48 hours. Cotreatment of cancer cells with RLI and Gem resulted in a substantial reduction in metabolic activity **(Fig. 1C)**, short-term cell viability and total live cell counts compared to either agent alone **(Fig. 1D)**. These short-term effects were further confirmed by long-term clonogenic assays, where the RLI and Gem treatment almost suppressed the ability of the cancer cells to form colonies **(Fig. 1E)**. The canonical viewpoint is that cell death by RLI or Gem is through apoptosis (12, 13). Immunoblotting revealed that RLI and Gem effectively reduced the phosphorylation of p38 (P-p38), indicating successful p38 MAPK inhibition. However, a marked decrease in activation of caspase-3 was observed in combination groups when compared with RLI or Gem alone **(Fig 1F and G)**. This suggests that while the combination treatment enhances apoptosis, it may be, at least in part, caspase independent. Further, we demonstrated that cotreatment with RLI and Gem showed a marked increase and in both early and late apoptosis **(Fig. 1H)**. Taken together, these data support that RLI enhances Gem-mediated apoptotic cell death in MIA PaCa-2 cells which may not involve canonical caspase activation. To validate our pharmacological findings of p38 MAPK inhibition, we performed genetic knockdown of p38 MAPK by siRNA (*sip38*). Immunoblot showed a marked decrease in both total and P-p38 levels when compared to non-target (*siNT*) counterparts **(Fig. 2A)**. *sip38* cells treated with Gem resulted in significant decrease in metabolic activity **(Fig. 2B)**, cell viability **(Fig. 2C)** and clonogenic potential **(Fig. 2D)**. Caspase-3 activation was reduced in Gem treated *sip38* cells compared to *siNT* cells **(Fig. 2E)**. An increase in late apoptosis and no significant difference in early apoptosis was observed in *sip38* cells treated with Gem **(Fig. 2F and G)**. These data showed that knockdown of p38 in MIA PaCa-2 cells phenocopied the effects of RLI, which warrants further investigation.

**Figure 1.**
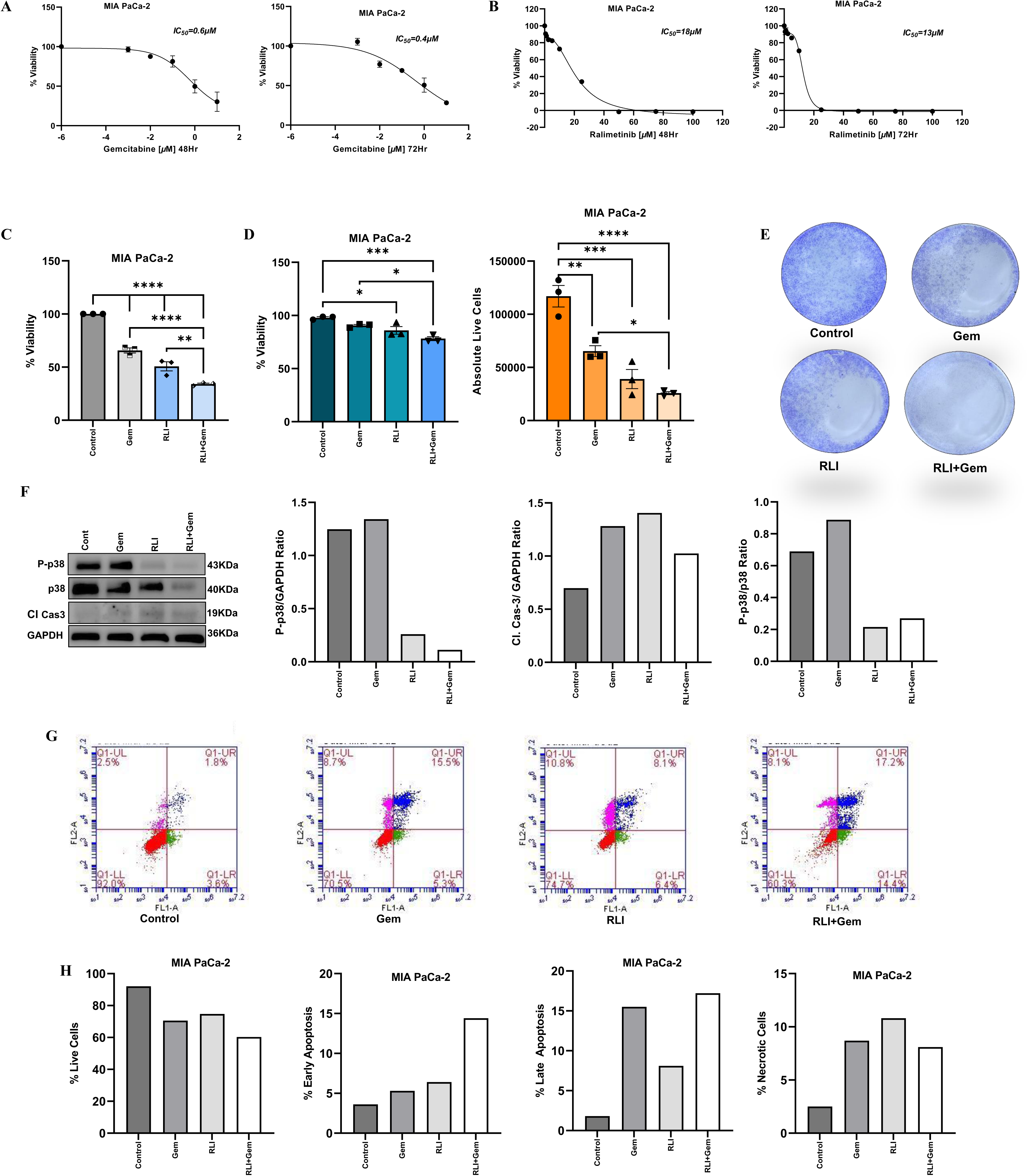
Ralimetinib (RLI) potentiates Gemcitabine (Gem)-induced cell death in MIA PaCa-2 cells. Dose-response curve for Gem **(A)** and RLI **(B)** in MIA PaCa-2 cells at 48 and 72 hours by MTT assay. Cells were pre-treated with RLI 13 µM for 24 hours followed by co-treatment with Gem 0.4 µM treatment for 48 hours. **(C)** Percentage of cell viability was measured using the MTT assay. **(D)** Cell viability and total live cells were determined by the Trypan Blue assay. **(E)** Crystal violet-stained colonies from cells treated with DMSO control, Gem (40nM), RLI (1.3uM), and their combination. **(F)** Whole cell lysates from DMSO control, Gem, RLI, and their combination were prepared. Immunoblot analyses were performed with indicated antibodies. Graph showing relative protein/protein ratios of P-p38/GAPDH, Cl. Cas-3/GAPDH, and P-p38/p38 (from left to right). **(G)** Apoptosis analysis by flow cytometry is shown for each group: Control, Gem, RLI, RLI+Gem (from left to right). The quadrants represent: Q1-LL (lower-left) for viable cells (Annexin V−/PI−), Q1-LR (lower-right) for early apoptotic cells (Annexin V+/PI−), Q1-UR (upper-right) for late apoptotic cells (Annexin V+/PI+), and Q1-UL (upper-left) for necrotic cells (Annexin V−/PI+). **(H)** Bar graphs showing quantitative summary of Live Cells, Early Apoptosis, Late Apoptosis, and Necrotic Cells for each treatment group. Data graphs represent mean ± SEM, n = 3 independent biological repeats. *P ≤ 0.05; **P ≤ 0.01; ***P ≤ 0.001; ****P ≤ 0.0001. ANOVA test was used when more than 2 groups were compared.

**Figure 2.**
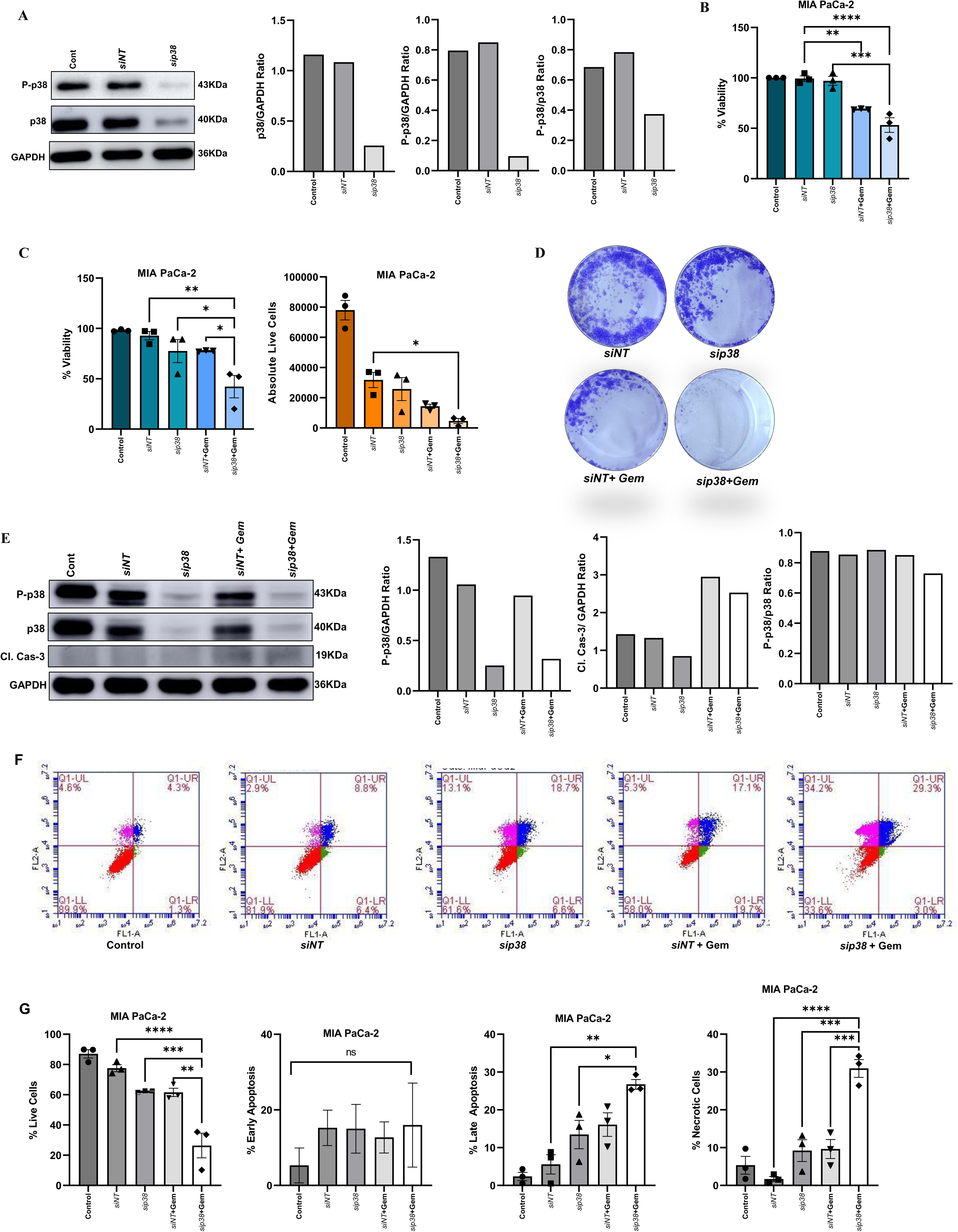
Genetic knockdown of p38 MAPK enhances Gem-induced cell death. MIA PaCa-2 cells were transfected with non-target control siRNA (*siNT*) or p38 siRNA (*sip38*) for 24 hours followed by Gem (0.4 µM) treatment for 48 hours. **(A)** Immunoblotting of P-p38, p-38 and GAPDH in the whole-cell lysates of MIA PaCa-2 cells.. Graph showing relative protein/protein ratios of p38/GAPDH, P-p38/GAPDH, and P-p38/p38 (from left to right).**(B)** Percentage of cell viability was measured using the MTT assay. **(C)** Percentage of viable cells and total live cells were determined by Trypan Blue assay. (D) Seven days colony formation assay with Gem (40 nM) in si*NT* or si*p38* targeted cells. **(E)** Immunoblots for indicated proteins in whole cell lysates of indicated. Graph showing relative protein/protein ratios of P-p38/GAPDH, Cl. Cas-3/GAPDH, and P-p38/p38 (from left to right). **(F)** Apoptosis analysis by Flow cytometry is shown for each group: Control, *siNT*, *sip38*, *siNT*+Gem, *sip38*+Gem (from left to right). **(G)** Bar graphs showing quantitative summary of Live Cells, Early Apoptosis, Late Apoptosis, and Necrotic Cells for each group. Data graphs represents mean ± SEM, n = 3 independent biological repeats. \**P* ≤ 0.05; \*\**P* ≤ 0.01; \*\*\**P* ≤ 0.001; \*\*\*\**P* ≤ 0.0001; ns non-significant. ANOVA test was used when more than 2 groups were compared.

### p38 MAPK targeting sensitizes Panc-1 cancer cells also for chemotherapy

We extended our findings to Panc-1, a KRAS mutant pancreatic cancer cell line. The IC₅₀ of Gem were 0.6 µM (48 hours) and 0.16 µM (72 hours) **(Fig. 3A)**, while that of RLI was 22 µM (48 hours) and 19 µM (72 hours) **(Fig. 3B)**, respectively. For combination studies, 19 µM RLI and 0.4 µM Gem were used. Pre-treatment with RLI (24 hours) followed by Gem (48 hours) significantly reduced metabolic activity **(Fig. 3C)**, viability, and live cell counts compared to single agents **(Fig. 3D)**. Immunoblotting confirmed inhibition of p38 MAPK (reduced P-p38) and showed diminished caspase-3 activation, suggesting partly caspase-independent apoptosis **(Fig. 3E)**. Cotreatment increased late apoptosis with a reduction in early apoptosis **(Fig. 3F and G)**. Genetic knockdown of p38 (*sip38*) **(Fig. 3H)** further validated these results, as *sip38* cells treated with Gem showed reduced metabolic activity **(Fig. 3I)**. Collectively, these data indicate that RLI enhances Gem-induced apoptosis in Panc-1 cells, potentially through caspase-independent pathways.

**Figure 3.**
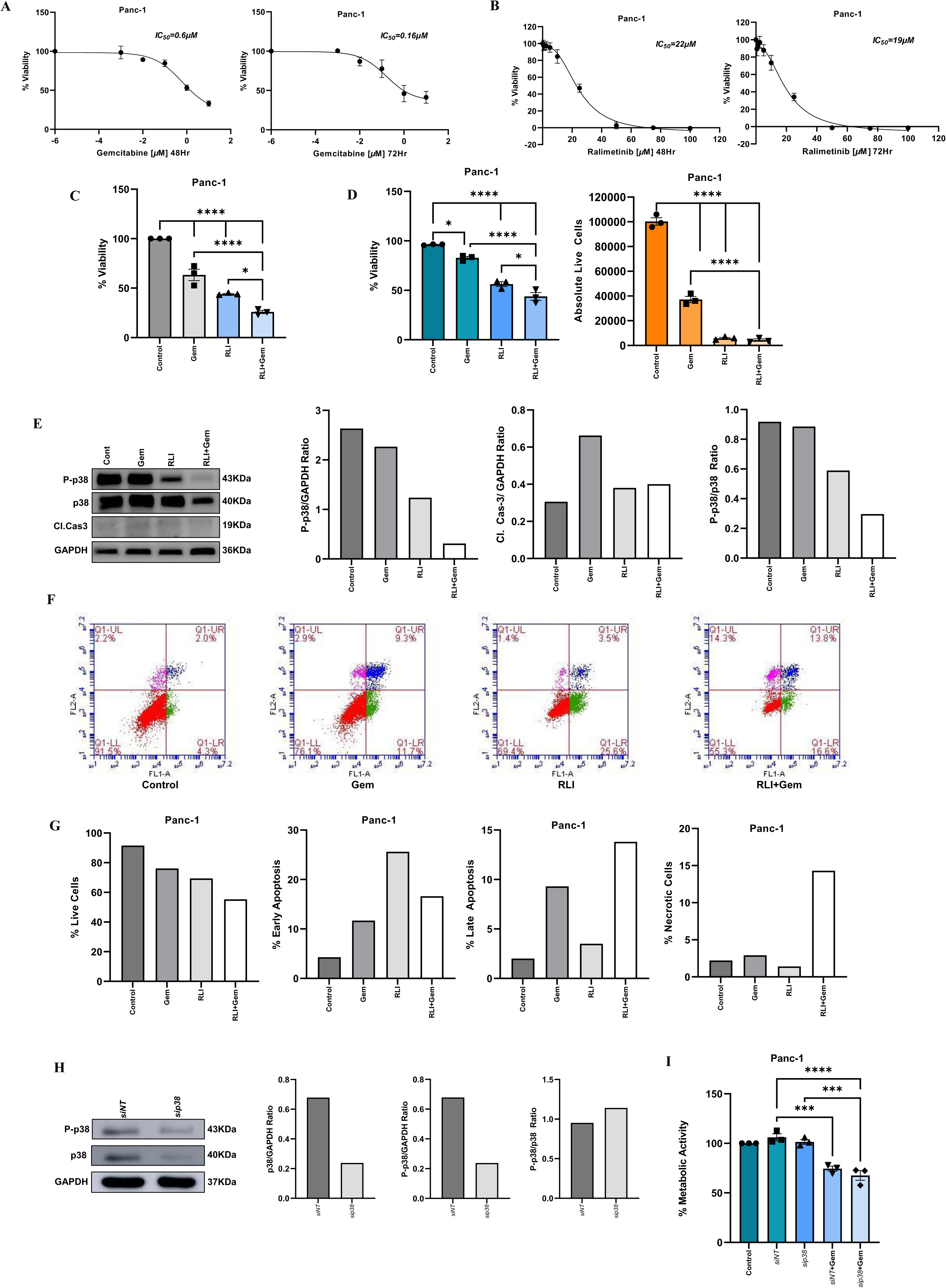
p38 MAPK Inhibition Sensitizes Panc-1 cells to Gemcitabine. Dose-response curve for Gem **(A)** and RLI **(B)** in Panc-1 cells at 48 and 72 hours by MTT assay. Cells were pre-treated with RLI (19 µM) for 24 hours followed by co-treatment with Gem (0.4 µM) treatment for 48 hours. **(C)** Percentage of metabolic activity, **(D)** cell viability and total live cells were determined by MTT and Trypan Blue assay respectively. **(E)** Whole cell lysates from DMSO control, Gem, RLI, and their combination were prepared. Immunoblot analyses were performed with indicated antibodies. Graph showing relative protein/protein ratios of P-p38/GAPDH, Cl. Cas-3/GAPDH, and P-p38/p38 (from left to right). **(F)** Apoptosis analysis by Flow cytometry and **(G)** quantification are shown for each indicated group. **(H)** Panc-1 cells were transfected with non-target control siRNA (*siNT*) or specific p38 siRNA (*sip38*) for 24 hours and whole-cell lysates were immunoblotted with indicated antibodies. Graph showing relative protein/protein ratios of p38/GAPDH, P-p38/GAPDH, and P-p38/p38 (from left to right). **(I)** Percentage of metabolic activity in *siNT* or *sip38* cells treated with or without Gem (0.4 µM) was measured using the MTT assay. Data graphs represent mean ± SEM, n = 3 independent biological repeats. \**P* ≤ 0.05; \*\**P* ≤ 0.01; \*\*\**P* ≤ 0.001; \*\*\*\**P* ≤ 0.0001. ANOVA test was used when more than 2 groups were compared.

### MAPK14 (p38α) is associated with poor prognosis in pancreatic cancer-A meta data analysis

To establish the role of p38MAPK as a prognostic marker for cancer progression, a comprehensive expression profile of all isoforms of p38MAPK was analyzed across pan-cancer tissues using Clinical Proteomic Tumor Analysis Consortium (CPTAC) datasets. A marked upregulation of MAPK14 (p38α) was seen in several malignancies, including pancreatic adenocarcinoma, while downregulation is observed in lung cancer **(Fig 4A)**. MAPK14 was significantly upregulated in tumor tissues (n=137) when compared to normal tissues (n=74) **(Fig 4B)**. High expression of MAPK14 showed an insignificant yet impactful trend towards poorer overall survival in patients **(Fig 4C)**. Elevated levels of MAPK14 emerged as an independent prognostic factor with worse survival outcomes in patients (Hazard Ratio = 4.59, p = 0.0059) **(Fig 4D)**. Age, sex, tumor stage, residual tumor status, and lymphovascular invasion were not independently associated with survival. Although stage IV disease and microscopic residual tumor (R1) exhibited a trend toward higher hazard, these did not reach statistical significance.

**Figure 4.**
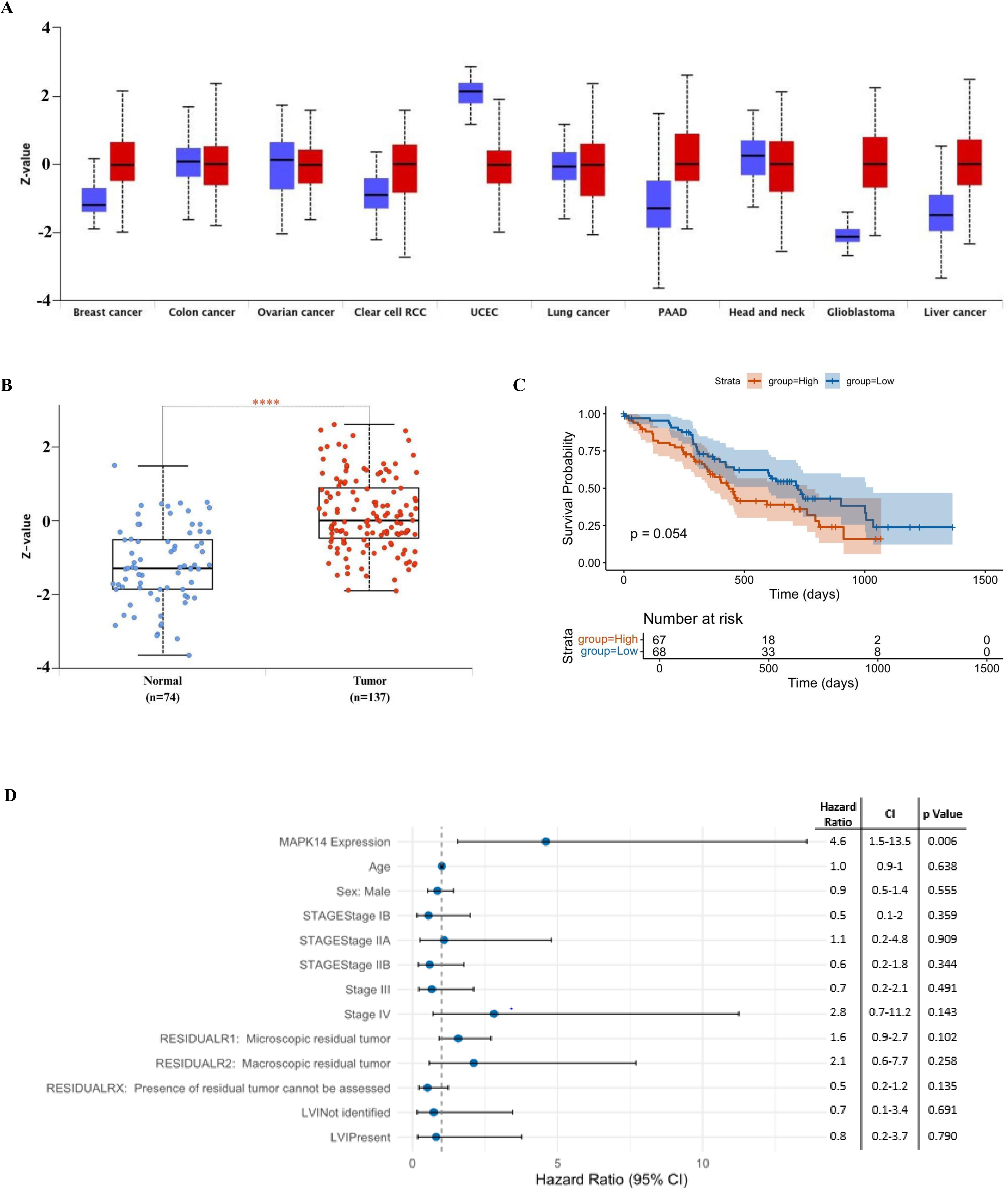
MAPK14 is overexpressed in Pancreatic Cancer and Associated with Poor Survival. **(A)** MAPK14 expression across multiple cancers. Boxplots show expression in normal (blue) vs tumor samples (red) from the CPTAC datasets. **(B)** Boxplot comparing MAPK14 expression between normal and PAAD tumor samples. **(C)** Kaplan–Meier survival analysis for patients stratified into high- and low-MAPK14 groups. **(D)** Multivariate Cox proportional hazards analysis (forest plot) displaying hazard ratios (HRs), 95% confidence intervals (CIs) and p-values for MAPK14 expression. The model was adjusted for [age, sex, stage, residual tumor, LVI]. A vertical dashed line indicates HR = 1.0; variables with p < 0.05 are denoted by an asterisk.

MAPK13 (p38δ) expression was significantly downregulated in PAAD patients suggesting MAPK13 does not play a crucial role in pancreatic cancer **(Supplementary Fig 1A and Supplementary Fig 1B)**. No significant association between MAPK13 expression level and patient survival (p=0.99) was observed **(Supplementary Fig 1C)**. Multivariate cox regression indicates high MAPK13 expression associated with reduced HR with a favorable prognostic impact **(Supplementary Fig 1D).** MAPK12 (p38γ) expression was significantly increased in PAAD tumor tissues **(Supplementary Fig 2A and Supplementary Fig 2B)** with decreased survival in patients with elevated expression of MAPK12 **(Supplementary Fig 2C)**. However, a protective effect on patient survival was observed which is associated with reduced hazard ratios and MAPK12 expression **(Supplementary Fig 2D).** MAPK11 (p38β) expressions were found insignificant in both normal and tumor tissues **(Supplementary Fig 3A and Supplementary Fig 3B)** and did not have a significant impact on overall patient survival **(Supplementary Fig 3C)** with decreased hazard ratios **(Supplementary Fig 3D)**. However, both MAPK12 and MAPK11 indicate poor prognosis with higher hazard ratios in advanced disease stages. Other clinical covariates like age, sex and lymphovascular invasion were indispensable on survival outcomes **(Supplementary Fig 2D and Supplementary Fig 3D)**. This suggests that high expression of MAPK14 is associated with poor survival outcomes in pancreatic cancer patients.

### MAPK14 Regulates Pro-Tumorigenic Pathways and Immune Infiltration in Pancreatic Cancer

To understand the biological role of MAPK14 in pancreatic cancer, we performed Gene Ontology (GO) enrichment analysis. Results revealed that MAPK14 is primarily involved in biological pathways (BP) related to stress-activated protein kinase cascades, cellular responses to UV light, and phosphorylation. Its molecular function (MF) is highly associated with kinase activity, particularly protein serine/threonine kinase activity, which is crucial for signal transduction **(Fig 5A-C)**.KEGG pathway analysis showed a significant and concomitant positive correlation between MAPK14 and Vascular Endothelial Growth Factor-A (VEGFA) signaling pathway **(Fig 5D and E)**. Given this strong correlation, we hypothesized that the combined expression of these genes could be a prognostic marker. Kaplan-Meier survival analysis showed that patients with both high MAPK14 and high VEGFA expression had a significantly worse survival than patients with both low MAPK14 and VEGFA groups **(Fig 5F)**. A significantly reduced overall survival in patients is associated with both high expression of MAPK14 and Hypoxia Inducible Factor-1 alpha (HIF1A), a key regulator of VEGFA **(Fig 5G)**. String analysis showed that MAPK14 is phosphorylated by upstream kinases like (Mitogen-activated protein kinase kinase) MAP2K3 and MAP2K6, that activate downstream kinases like Mitogen-activated protein kinase-activated protein kinase MAPKAPK2 and MAPKAPK3 **(Fig 5H).** MAPK14 modulates the transcriptional activity of (Activating transcription factor-2) ATF2 (14, 15) and (Jun proto-oncogene, AP-1 transcription factor subunit) JUN that promotes tumor growth and metastasis (16, 17) **(Fig 5H)**. In addition, MAPK14 interacts with (Mitogen-activated protein kinase-1) MAPK1 (ERK2, a key component of classical ERK pathway) that supports tumor resistance. Interaction of MAPK14 with tumor suppressor (Tumor protein 53) TP523 is associated with cell cycle arrest and apoptosis **(Fig 5H)**. These results showed that MAPK14 plays a crucial role in exerting pro-tumorigenic effects in pancreatic cancer. Furthermore, spearman correlation analysis between MAPK14 expression and different types of immune cell infiltration in 179 PAAD samples were carried out. Scatter plots with positive correlation lines (rho values between 0.2 – 0.4) were observed for CD8^+^ T cells, Th1 cells and NK cells showing significant anti-tumor potential of MAPK14 expression **(Fig 6)**. In parallel, Tregs and macrophages showed weaker yet significant positive correlation with high MAPK14 expression **(Fig 6)**. This result indicates that higher MAPK14 expression leads to greater infiltration of immune cells in pancreatic cancer.

**Figure 5.**
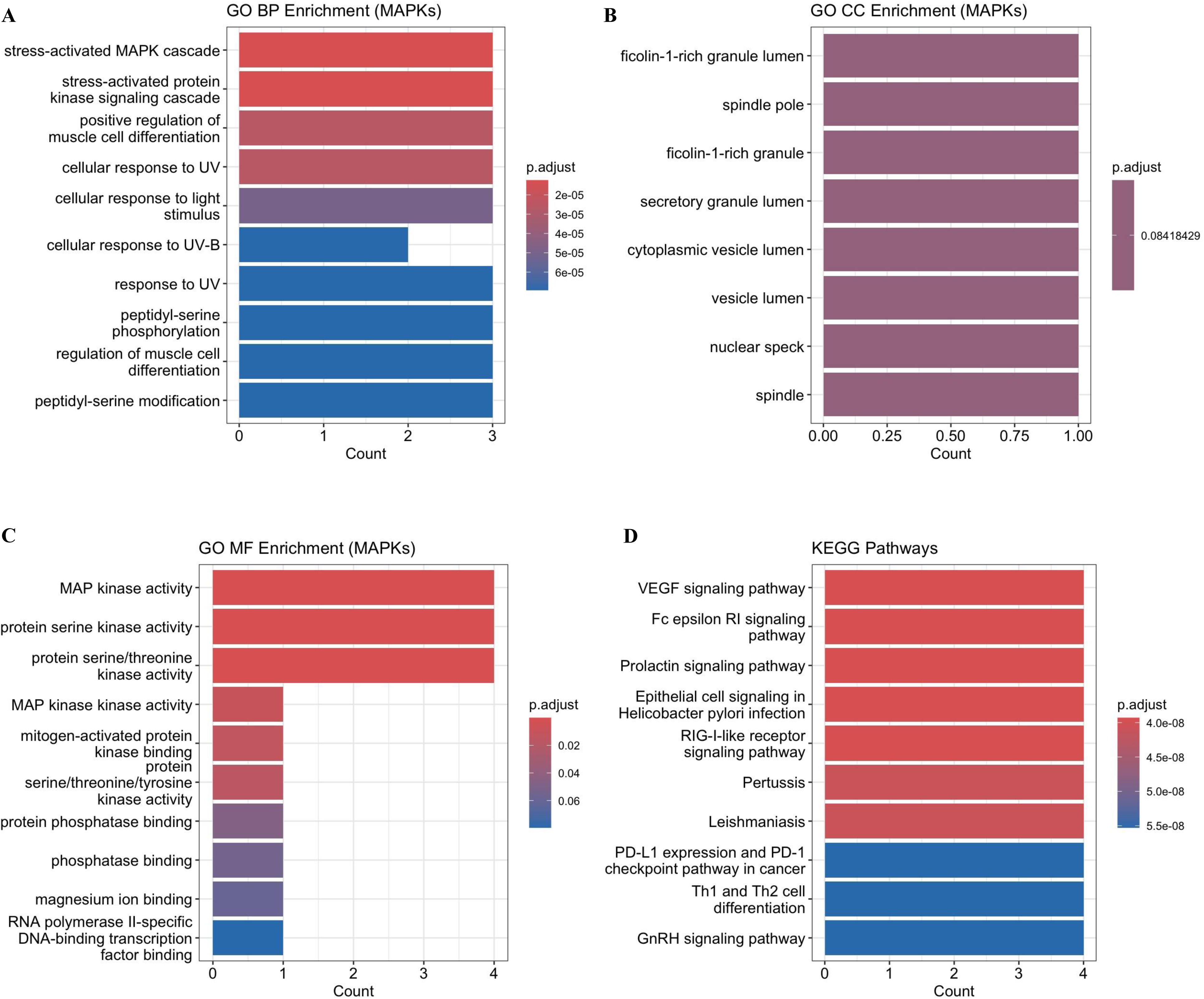
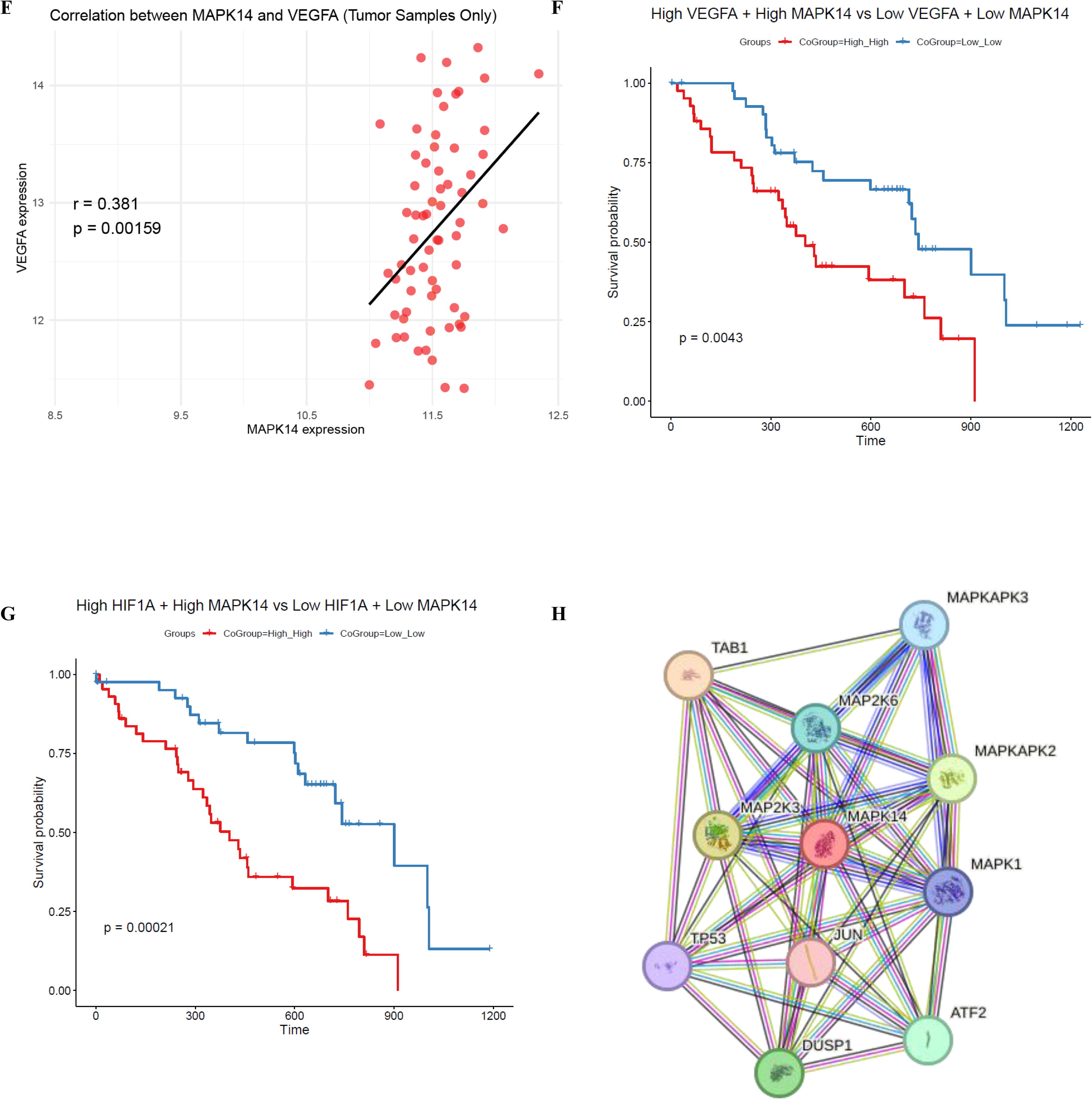
Integrative Analysis of MAPK14: Functional enrichment, protein-protein interaction, and correlation with VEGFA (A-D) Functional enrichment analysis of MAPKs. Gene Ontology (GO) and KEGG pathway enrichment analyses were performed for p38 MAPK. **(A)** GO Biological Process (BP) enrichment revealed significant enrichment in stress-activated MAPK cascade, cellular responses to UV/light stimulus, and regulation of muscle cell differentiation. **(B)** GO Cellular Component (CC) enrichment indicated associations with vesicle lumen, spindle pole, and nuclear speck, although these terms were less significant. **(C)** GO Molecular Function (MF) enrichment showed MAP kinase activity, protein serine/threonine kinase activity, and kinase binding as the most enriched functions. **(D)** KEGG pathway analysis showed highly enriched terms in VEGF signaling, Fc epsilon RI signaling, PD-L1/PD-1 checkpoint pathway in cancer, and Th1/Th2 differentiation. **(E)** Pearson correlation analysis identified a moderate but significant positive correlation between VEGFA and MAPK14 expression (r = 0.313, p = 0.000169), indicating association between MAPK14 activity and angiogenic signaling. **(F-G)** Kaplan–Meier survival analysis revealed patients with high VEGFA + high MAPK14 expression and high HIF1A + high MAPK14 expression had significantly worse survival outcome than low expression groups. **(H)** String analysis revealed protein– protein interaction (PPI) network analysis of MAPK14 and its contact with key regulators such as, ATF2, TP53, JUN, and DUSP1 highlighting its central importance in MAPK signaling and stress responses. **( I )** Spearman correlation analysis was performed to evaluate the association between MAPK14 expression and immune cell subsets in 179 PAAD samples.

**Figure 6.**
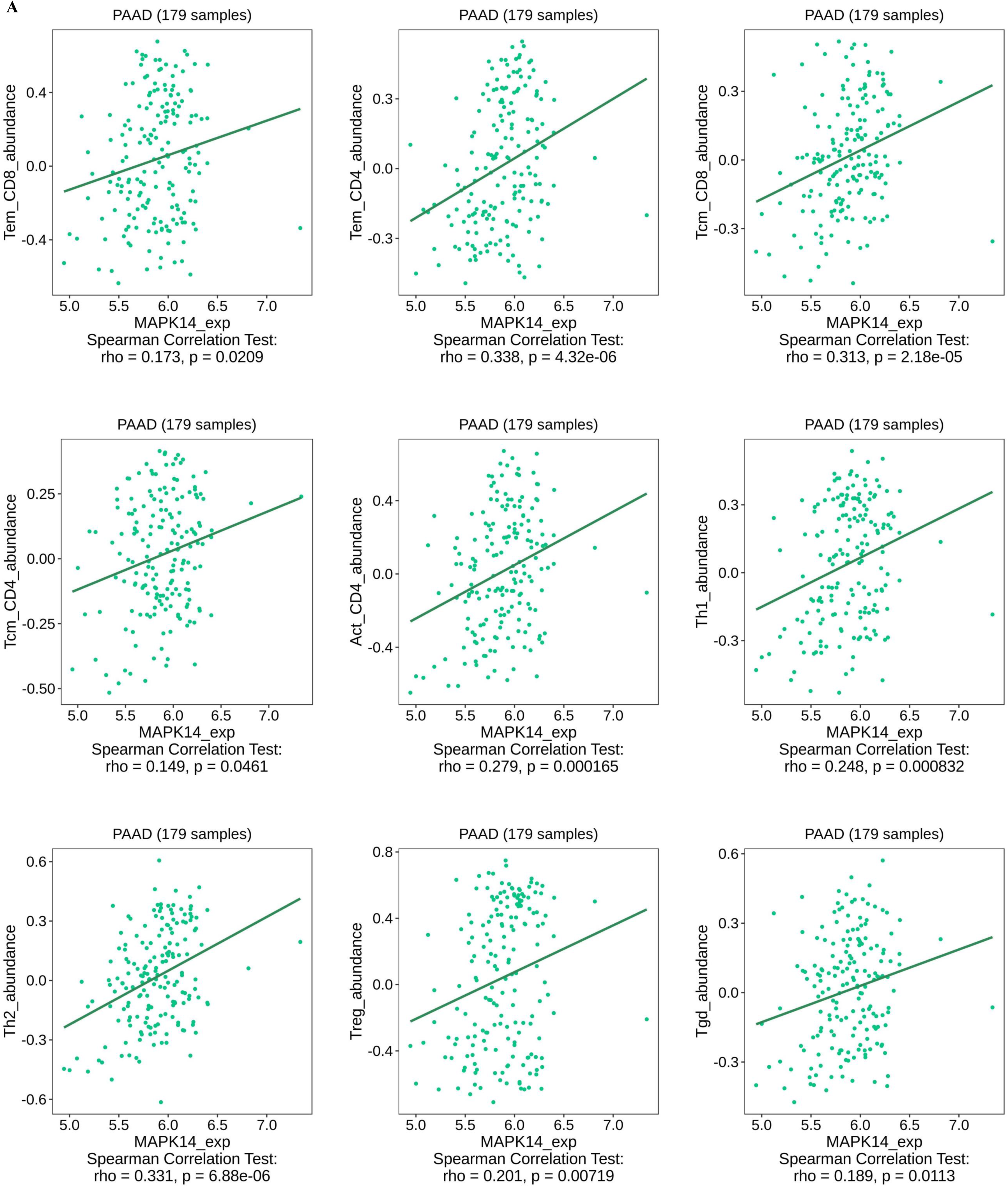
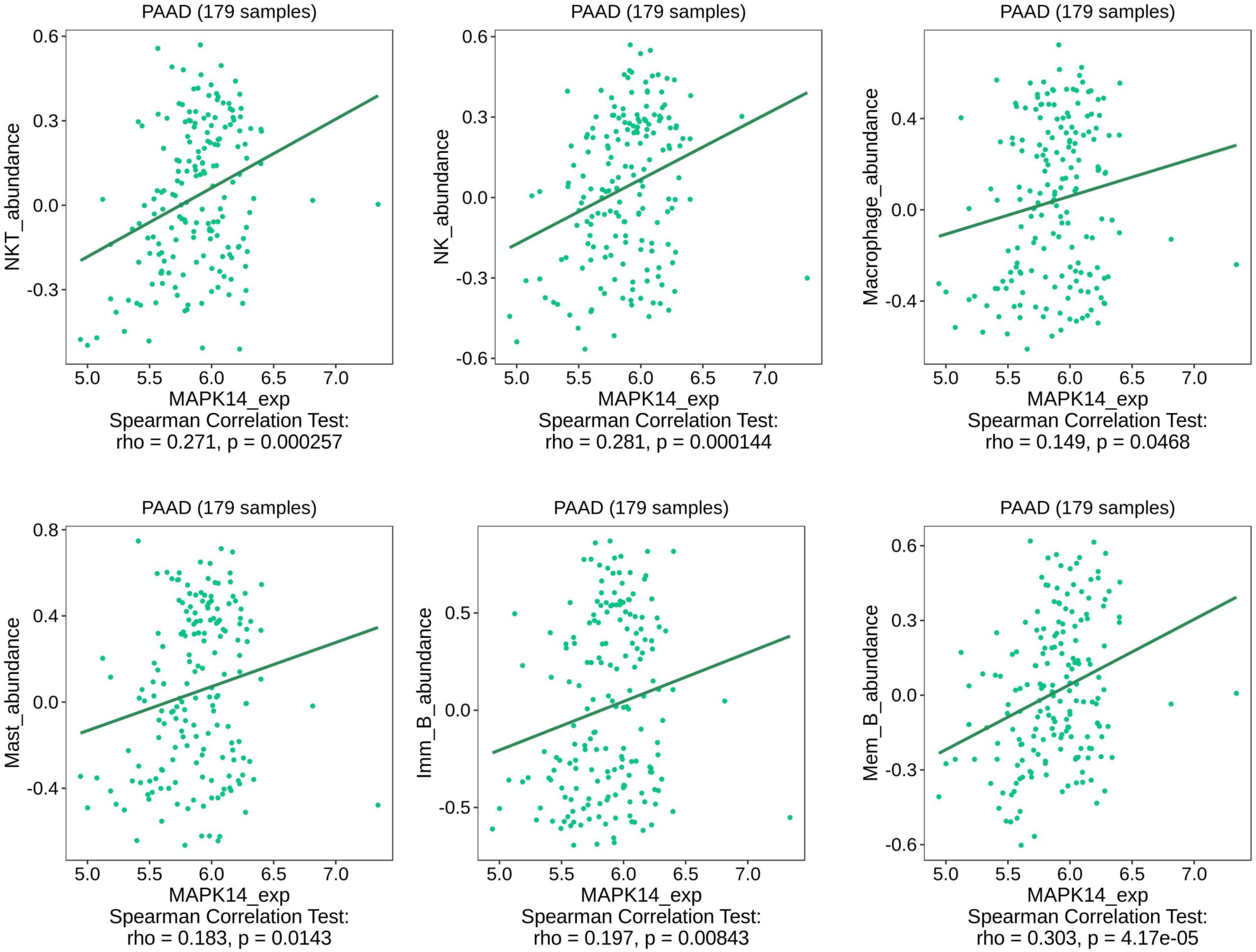

## Discussion

Despite of improvements in diagnostic and therapeutic modalities, pancreatic cancer is still one of the deadliest cancers with a poor prognosis having less than 11% five-year median survival rate. It is the 6^th^ leading cause of cancer related mortalities worldwide. 90% of pancreatic cancer patients are manifested as pancreatic ductal adenocarcinoma (PDAC). This is manifested with the accumulation of various angiogenic factor which renders tumor endothelium impermissive for T cell infiltration to support cancer progression from chronic pancreatitis which render pancreatic tumors refractory for various treatment modalities. Since macrophage infiltration and their *in-situ* polarization is decisive for neoplastic transformation of inflamed pancreas. Because of this targeted activation of the tumor microenvironment may overcome endothelium barrier dysfunctions / anergy which is a major limitation of existing cancer immunotherapy. Out of various signaling component which remains hyperactive in tumors, MAPK family members like extracellular signal-regulated kinase (ERK), c-Jun N-terminal kinase (JNK), and p38 are over expressed in variety of tumors. These proteins promote in situ polarization of tumor infiltrating M1 tumor associated macrophages (TAM) towards M2 TAM which confer poor prognosis in cancer patients. A study done by Prakash *et al.* 2016 on low-dose gamma irradiation has shown tumor rejection in mice model of insulinoma by causing induction of M1 associated effector cytokines through down regulation of p38 MAPK. Therefore, we hypothesize inhibiting p38 MAPK will confer protective immunity against pancreatic cancer and potentiate the efficacy of existing anti-tumor therapies. P38MAPK is downstream of VEGFR2 and regulate set of miRNA which are involved in the angiogenesis. KRAS mutation triggers MAPK mediated inflammation and desmoplastic stroma which play a decisive role for the growth of pancreatic cancer. Additionally, inhibiting p38 MAPK, alone or in combination with chemotherapy will retune M2 TAM towards M1 phenotype, enhance tumor antigen presentation by M1 repolarized TAM to T cells and their subsequent reactivation. This is believed to convert “cold” tumor into “hot” and permeable tumors and tweak immune mediated rejection of PDAC. In view of this, our approach is quite logical and has translational potential for improving the response of PDAC patients for various cancers directed interventions

## Conclusion

In this study we propose to inhibit MAPK pathways in PDAC cells for recalibrating their sensitivity toward chemotherapy only but due to involvement of p38MAPK in EFGR / IGF/VEGFR signalling network we believe that introduction of p38MAPK based approach would improve the response of PDAC towards various immunotherapies targeting these growth factor associate pathways. P38MAPK based approach is also believed to enhance the sensitivity of various Immune check point blockade therapies which are targeting T cells responses. In view of this, we believe that our study not only perpose and also re-inforce the significance of p38MAPK based interventions as potential host directed approach for managing tumor burden in patients

## Materials and Methods

### Cell Culture

Panc-1 and MiaPaCa-2 human pancreatic cancer cell lines were obtained from National Center for Cell Sciences (NCCS), Pune, India. Cells were maintained at 37° C in a humidified incubator at 5% CO_2_ and were grown in DMEM medium (Gibco), supplemented with 10% fetal bovine serum (FBS, Gibco), and 1X antibiotic-antimycotic (Gibco).

### Cell Viability Assay

Cell viability was assessed using the MTT assay (SRL) and Trypan Blue (Gibco). 5,000– 7,000 cells per well optimized for each cell line were seeded into 96-well plates on Day 1. Drug treatments were administered on Day 2, and viability was measured on Day 3 or Day 4. For MTT assay, MTT reagent was added and incubated for 3–4 hours, followed by solubilization of formazan crystals. Absorbance was recorded at 570 nm using a microplate reader. Dose-response curves and IC₅₀ values were generated using GraphPad Prism 9. For Trypan Blue assay, cells were trypsinized and diluted in 0.4% trypan blue solution in the ration of 1:1, loaded in haemocytometer and counted.

### Colony Forming Assay

7,500 cells per well were seeded into a 12-well plate and were treated next day, cells were kept under treatment for a total of 7 days. Colonies formed in each well were washed with DPBS (Himedia), fixed with cold methanol and kept at -20°C for 20 minutes, and stained with 0.5% Crystal violet solution (Millipore Sigma).

### Western Blotting

Total protein was extracted using RIPA lysis buffer (Sigma) with protease inhibitor (G-Biosciences) and phosphatase inhibitor (Roche). Protein concentrations were determined using the Pierce Rapid Gold BCA Protein Assay Kit (Thermo Scientific). Equal amounts of protein were resolved on Tris-Glycine SDS-PAGE gels and transferred onto polyvinylidene difluoride (PVDF) membrane (Millipore). Membranes were blocked, incubated with primary and HRP-conjugated secondary antibodies, and visualized using the femtoLUCENT Plus HRP ECL detection system (G-Biosciences).

### Apoptosis Assay

Apoptosis induction was measured using Annexin V-FITC apoptosis kit (Invitrogen eBioscience). MIA PaCa-2 and Panc-1 cells were seeded in a 6-well plate. Cells treated with DMSO control, Gem, RLI and their combination for 72Hr. Cells were trypsinized, washed with DPBS, and resuspended in 195µl 1X binding buffer. 5µl Annexin V-FITC was added to the cell suspension and incubated for 10min at room temperature. Cells were washed and resuspended in 190µl 1X binding buffer. 10µl Propidium Iodide was added to the cell suspension and FACS analysis performed.

### siRNA Transfection

Genetic knockdown studies for human MAPK14 were performed in MIA PaCa-2 and Panc-1 cell lines. p38α MAPK14 (sc-29433) siRNA was purchased from Santa Cruz Biotechnology, consisting of 4 target-specific 19-25 nt siRNAs.

### In silico analysis Data Acquisition

Expression data for MAPK isoforms (MAPK11, MAPK12, MAPK13, and MAPK14) in pancreatic adenocarcinoma (PAAD) samples was obtained from the CPTAC dataset through the cBioPortal Data Hub. Processed RNA-seq data and clinical data were downloaded using the cBioPortalData R package.

### Differential Expression Analysis

Using the UALCAN web portal (https://ualcan.path.uab.edu/analysis-prot.html), differential gene expression analysis was performed through CPTAC-derived datasets. Gene expression levels of MAPK11, MAPK12, MAPK13, and MAPK14 were analyzed pan-cancer and in pancreatic adenocarcinoma (PAAD), comparing tumor and normal samples.

### Survival and Prognostic Analysis

For assessing the prognostic value of MAPKs, Kaplan-Meier survival analysis and Cox regression models were performed. Kaplan-Meier analysis was conducting using the survival and survminer R packages. Patients were divided into high- and low-expression groups according to the median expression of each gene. Log-rank test was used to evaluate the survival differences. Cox proportional hazards regression was performed in both univariate and multivariate settings to assess the independent prognostic significance of MAPK expression, by adjusting relevant clinico-pathological variables including age, sex, tumor stage, residual tumor, and lymphovascular invasion. Hazard ratios (HR) with 95% confidence intervals (CI) were reported.

### Co-expression Survival Analysis

MAPK14 and VEGFA were analyzed jointly to evaluate the prognostic relevance of their co-expression. Samples were grouped into 4 categories on the basis of high and low expression combinations. Kaplan-Meier survival analysis was performed on these categories, and survival curves were compared using the log-rank test.

### Functional Enrichment Analysis

For determining the structurally and functionally associated proteins, protein-protein interaction (PPI) networks were constructed using the STRING database (https://string-db.org/) for MAPK14. Further functional enrichment analyses were performed using the cluster Profiler R package. Kyoto Encyclopedia of Genes and Genomes (KEGG) pathway and Gene Ontology (GO) enrichment analyses (biological process, cellular component, and molecular function) were performed. Barplots were generated to visualize the results using the ggplot2 R package.

### Immune Infiltration Analysis

TISDIB database (http://cis.hku.hk/TISIDB/) was utilized to retrieve the immune cell infiltration correlations with MAPK14 expression. Spearman’s correlation was used to evaluate the relationship between gene expression and immune cell infiltration levels.

### Statistical Analysis

Data analysis was performed using GraphPad Prism software version 9.00 for Windows. Differences between two groups were performed using Student’s unpaired, 2-tailed t test and for more than two groups using one-way ANOVA. A p-value < 0.05 was considered statistically significant. Data are shown as the mean ± SEM.

## Acknowledgment

This work was supported by a grant from ICMR grant to HP. Authors thanks the excellent technical support which they received from Central Flow Cytometry facility of university

## Conflict of Interest

Author declare that this study was conducted without any conflict of interest

## Author contribution

**VM** conducted cellular experiments, **KG & GS** acquired and analysed TCGA / in silico data, **MB** supervised the experiments and wrote manuscript, **CG** and **PD** contributed with the technical resources, **MG** provided tool for the analysis **HP** conceptualized idea, supervised entire study and edited manuscript

## Legends to Supplementary figures

**Suppl Figure 1.**
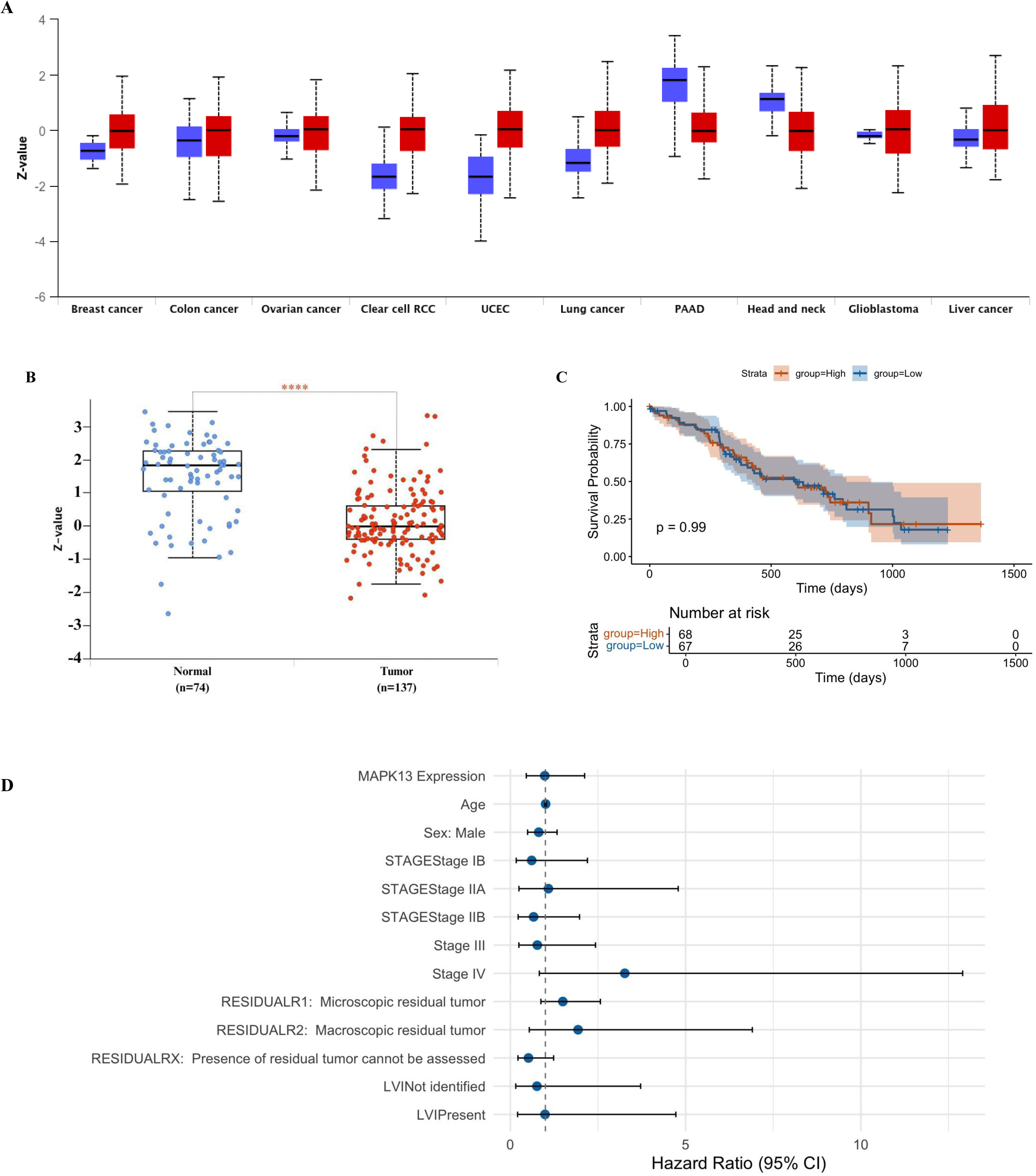
Protein expression and prognostic significance of MAPK13 in Pancreatic Cancer patients. **(A)** MAPK13 expression across multiple cancers. Boxplots show protein expression in normal (blue) vs tumor samples (red) from the CPTAC datasets. **(B)** Boxplot comparing MAPK13 expression between normal and PAAD tumor samples. **(C)** Kaplan–Meier survival analysis for patients stratified into high- and low-MAPK13 groups. **(D)** Multivariate Cox proportional hazards analysis (forest plot) displaying hazard ratios (HRs), 95% confidence intervals (Cis). The model was adjusted for [age, sex, stage, residual tumor, LVI]. A vertical dashed line indicates HR = 1.0; variables with p < 0.05 are denoted by an asterisk.

**Suppl Figure 2.**
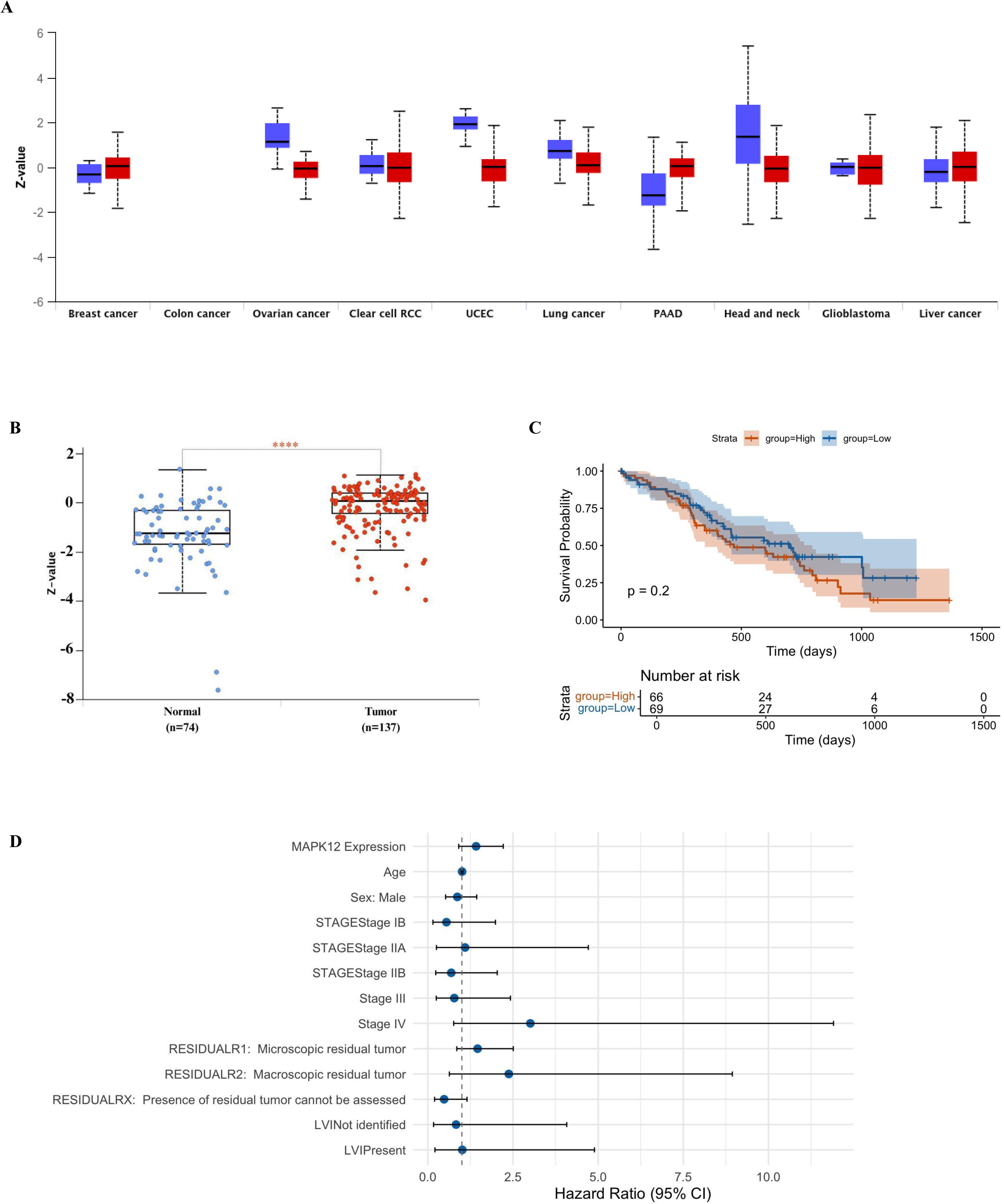
Protein expression and prognostic significance of MAPK12 in Pancreatic Cancer patients. **(A)** MAPK12 expression across multiple cancers. Boxplots show protein expression in normal (blue) vs tumor samples (red) from the CPTAC datasets. **(B)** Boxplot comparing MAPK12 expression between normal and PAAD tumor samples. **(C)** Kaplan–Meier survival analysis for patients stratified into high- and low-MAPK12 groups. **(D)** Multivariate Cox proportional hazards analysis (forest plot) displaying hazard ratios (HRs), 95% confidence intervals (Cis). The model was adjusted for [age, sex, stage, residual tumor, LVI]. A vertical dashed line indicates HR = 1.0; variables with p < 0.05 are denoted by an asterisk.

**Suppl Figure 3.**
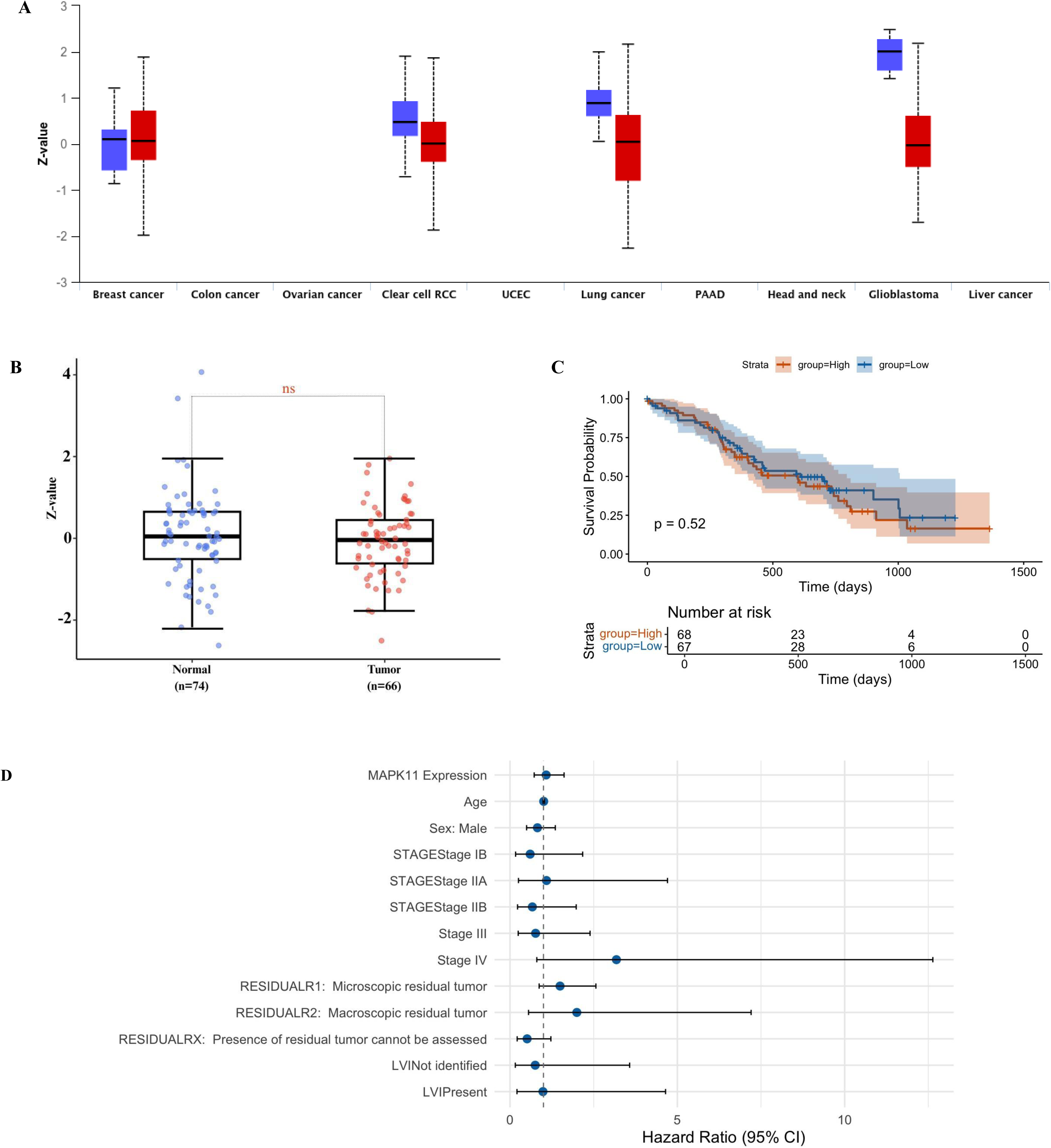
Protein expression and prognostic significance of MAPK11 in Pancreatic Cancer patients. **(A)** MAPK11 expression across multiple cancers. Boxplots show protein expression in normal (blue) vs tumor samples (red) from the CPTAC datasets. **(B)** Boxplot comparing MAPK11 expression between normal and PAAD tumor samples. **(C)** Kaplan–Meier survival analysis for patients stratified into high- and low-MAPK11 groups. **(D)** Multivariate Cox proportional hazards analysis (forest plot) displaying hazard ratios (HRs), 95% confidence intervals (Cis). The model was adjusted for [age, sex, stage, residual tumor, LVI]. A vertical dashed line indicates HR = 1.0; variables with p < 0.05 are denoted by an asterisk.

## Notes

### Competing Interest Statement

The authors have declared no competing interest.

